# Comparative genomics of a novel *Erwinia* species associated with the Highland midge (*Culicoides impunctatus*)

**DOI:** 10.1101/2023.09.28.559968

**Authors:** Jack Pilgrim

## Abstract

*Erwinia* (Enterobacterales: Erwiniaceae) are a group of cosmopolitan bacteria best known as the causative agents of various plant diseases. However, other species in the group have been found to play important roles as insect endosymbionts supplementing the diet of their hosts. Here, I describe *Candidatus* Erwinia impunctatus (Erwimp) associated with the Highland midge *Culicoides impunctatus* (Diptera: Ceratopogonidae), an abundant biting pest in the Scottish Highlands. The genome of this new *Erwinia* species was assembled using hybrid long and short read techniques, and a comparative analysis was undertaken with other members of the genus to understand its potential ecological niche and impact. Genome composition analysis revealed that Erwimp is similar to other endophytic and ectophytic species in the genus and is unlikely to be restricted to its insect host. Evidence for an additional plant host includes the presence of a carotenoid synthesis operon implicated as a virulence factor in plant-associated members in the sister genus *Pantoea*. Unique features of Erwimp include several copies of intimin-like proteins which, along with signs of genome pseudogenization and a loss of certain metabolic pathways, suggests an element of host restriction seen elsewhere in the genus. Furthermore, a screening of individuals over two field seasons revealed the absence of the bacteria in *C. impunctatus* in the second year indicating this microbe-insect interaction is likely to be transient. These data suggest that *Culicoides impunctatus* may have an important role to play beyond a biting nuisance, as an insect vector transmitting Erwimp alongside any conferred impacts to surrounding biota.

## 3. Introduction

Animals and plants forge varying interactions with bacteria ranging from pathogenic (antagonistic), through to commensal (neutral) and mutualistic (beneficial). Within these categories, insect-microbe interactions are of particular interest due to insect roles as crop pests, insect vectors and pollinators. Certain bacteria are able to persist within cells and specialized tissues of insects (endosymbiosis) which has led to several instances where the fitness of the insect is intrinsically linked with that of the bacteria (1). Such phenomena have led to a range of phenotypes such as host reproductive manipulation (2), defense against natural enemies (3,4) and diet supplementation (5,6). In other cases, bacteria have fleeting interactions with insects, either utilizing them as vectors on their mouthparts (7) and cuticle surface (8) or by carrying them as gut bacteria (9) where they can be passed on to plants, animals or other environments.

An exemplary group of bacteria which embodies this astonishing range of interfaces with invertebrates are *Erwinia* spp. (Enterobacterales: Erwiniaceae). Endosymbiotic examples include *Erwinia haradaeae* which has evolved as an obligate primary (essential) endosymbiont supplementing *Cinara* aphids with essential B-vitamins and has a reduced genome (∼1.1 Mbp) with a relative enrichment of house-keeping genes and those genes relevant for nutritional symbiosis (10). On the other hand, *Erwinia dacicola* present in the olive fly *Bactrocera oleae* has a more complex lifecycle as a facultative endosymbiont which transitions between an intracellular stage in larval midguts to an extracellular phase in the gut lumen of adults (11) where it is thought to supplement its host with nitrogen through a urease operon (12). Despite these specialised insect roles, a majority of *Erwinia* are phytopathogens which are in part horizontally transmitted by insects feeding and resting on a primary plant host (13) (14). The most well-studied plant pathogens include *Erwinia amylovora* and *Erwinia pyrifoliae* which can cause devastating disease outbreaks in members of the Rosaceae family, however, other species (e.g., *Erwinia tasmaniensis*) are considered epiphytic non-pathogens (15). More rarely, Erwinia have been isolated from fungi (16) and humans (17,18) where they infrequently cause disease in the latter.

In the absence of ecological and biochemical characterisation of newly identified bacteria, comparative genomics can be used to narrow down ecological niches as well as suggest the nature of interactions with a host. In this study, I use comparisons of known Erwinia genomes encompassing insect endosymbionts (unculturable) and free-living (culturable) members of the genus to investigate the metabolic potential, genome composition and virulence factors of a novel strain, Candidatus *Erwinia impunctatus* (Erwimp), identified in the pest species of biting midge, *Culicoides impunctatus* (Diptera: Ceratopogonidae). Present across most of Northern Europe, these small blood-feeding insects are particularly notable in the Western Scottish highlands where huge numbers form as a result their ability to reproduce successfully once without a blood-meal (autogeny) (19). Their sheer numbers, as well as multiplicity of environments (adults feed on both animals and plants), mean the nature of interaction between *C. impunctatus* and Erwimp could have impacts on the insect itself or the flora and fauna they interact with.

## 4. Methods

### Sample collection, DNA extraction, genome assembly and annotation

Five hundred and thirty-one live female *Culicoides impunctatus* were collected by aspiration from the author’s (JP) skin in Kinlochleven, Scotland (56° 42’ 50.7”N -4° 57’ 34.9”E) in September 2020. Four-hundred and eighty *C. impunctatus* individuals were pooled, while the other 51 were kept for individual screening for any subsequent bacteria identified through sequencing. These were then homogenised and high-molecular weight DNA was extracted using the method described in Davison et al. (20). Oxford Nanopore libraries were generated using the SQK-LSK109 Ligation Sequencing Kit and sequenced on a Minion R9.4.1 flow cell. Raw reads were base called using Guppy version 4.0.15 (Oxford Nanopore) and the high-accuracy (hac) option. All reads >500⍰bp in length which had an average phred (Q) score of above 10 were filtered using NanoFilt v2.7.180 (21). Assembly of these reads took place using Flye v2.8.181 with default options (22).

High quality short-read libraries were also generated and sequenced from the same DNA samples using a Kapa HyperPrep kit (Roche) and a DNBseq G50 platform by BGI Genomics (Hong Kong) to correct the initial assembly. Filtering of raw DNBseq reads was performed by BGI Genomics’ by removing adapters using SOAPnuke v2.1.482 (23) and keeping reads with a mean Q score of >20. Filtered reads were assembled using MEGAHIT v1.2.983 before binning contigs with MetaBAT 2 v2.12.184 (24). The identities of bins were checked with CheckM v1.1.378 (25) before reads were mapped back to the contigs identified using ‘perfect mode’ in BBMap v38.8785 (26) and filtered using SAMtools v1.1186 (27). These mapped reads were then used to polish SNPs and indels in the initial nanopore assembly using two rounds with Pilon v1.2387 and the --*fix bases* option (28). Completeness was assessed using BUSCO v5.1.2 (29) through the gVolante server (30) based on 366 single-copy bacterial markers for gammaproteobacteria (gammaproteobacteria odb10 database). Annotation of the final polished genome was undertaken using PROKKA v1.1388 (31) with identification of secondary metabolites using antiSMASH v7.0 (32).

### Phylogenomics and ANI

As the provisional taxonomic assignment for a ∼3.5Mbp circular contig was as a member of the Erwiniaceae (Supplementary File S1), a total of 48 genomes from this family spanning the genera *Erwinia, Pantoea, Tatumella, Izakhiella* and *Mixta* (as well as four outgroup genomes; *Pectobacterium brasiliense, Serratia marcescens, Klebsiella pneumoniae* and *Shigella dysenteriae*) were downloaded from NCBI for further phylogenomic assessment (Supplementary File S2). To identify and extract single-copy orthologues, anvi’o v7 (33) was used to create a pangenome database. One-hundred and forty-four single-copy orthologues with a minimum geometric identity of 90% (to remove poorly aligned genes) from all genomes were then filtered. The *anvi-get-sequences-for-gene-clusters* program was then used with the --concatenate-gene-clusters option to extract the 144 concatenated orthologues for each genome. Gblocks v0.91b (34) was then used to exclude areas of the alignment with excessive gaps or poor alignment. ModelFinder (35) then determined the LG+I+G4 model to be used after selection using the Bayesian information criteria. A maximum likelihood (ML) phylogeny was then estimated with IQTree (36) using an alignment of 38,623 amino acids and 1,000 ultrafast bootstraps. To supplement the phylogenomic inferences, an average nucleotide identity (ANI) was also ascertained using the *anvi-compute-genome-similarity* program and pyani (37).

### Clusters of Orthologous Genes (COG), pseudogene and metabolic profiling

To compare the genomic profiles of free-living (culturable) and endosymbiotic (unculturable) *Erwinia* with Erwimp, Clusters of Orthologous Genes (COG) profiles were created using the *anvi-run-ncbi-cogs* program before the proportions of each category were plotted as a bar plot in ggplot2 v3.4.2 (38) using Rstudio v2022.02.0 (39). Pseudofinder v1.1.0 (40) was used to identify the proportion of pseudogenes in all Erwinia genomes. To compare metabolic profiles between Erwinia genomes in the pangenome database, Anvi’o was used with the programs *anvi-run-kegg-kofams* and *anvi-estimate-metabolism*, which utilised previous annotation of genes with KEGG orthologs (KOs) (41). The outputted completeness matrix for amino acid, lipid, carbohydrate, and vitamin metabolic pathways was then inputted into pHeatmap v1.0.12 (42) and visualised using Rstudio. In addition, PhyloFlash v3.4 (43)was utilised to search for SSU rRNAs possibly associated as other sources of Erwimp within the metagenome.

### Virulence factors and plant gene identification

To gain a better understanding of the ecological impact of Erwimp, common virulence factors (44) and plant-symbiosis genes (45) identified previously were compiled and searched for using PathwayTools v26.5 (46). Presence/absence matrices were created for each of these pathways before being plotted using pHeatmap. Proksee (47) was then used to visualise the circular chromosome of Erwimp and ORFs found in PathwayTools were then annotated in Inkscape v1.1 (48). Clinker (49) plotted the similarity of carotenoid biosynthetic genes discovered using AntiSMASH. Interpro v95.0 (50) provided functional analysis by predicting domains for Intimin-like genes.

### Targeted Erwinia screening of *Culicoides impunctatus* individuals

To understand the distribution of Erwimp in *C. impunctatus* populations, a remaining 51 individuals from the 2020 (Kinlochleven) catch, alongside 64 individuals caught in 2021 in neighbouring areas to the initial catch, were stored in 75% ethanol at -20°C before individual DNA extracting as described previously (51). Screening of the *COI* gene (primers C1-J-1718/C1-N-2191) was initially assessed as a means of quality control of extracts by conventional PCR (52). DNA extracts which passed quality control were then screened using a conventional PCR with primers based on the DNA Gyrase subunit B (GyrB) gene (Erwimp_Gyrb_1832F: 5-ATCGTCCGGTATTGTCTGCG-3’; Erwimp_Gyrb_2194R 5’-ATCCACGACGAGACTCCTT-3’). PCR assays consisted of a total of 15 μL per well, comprising of 7.5 μL GoTaq® Hot Start Polymerase, 5.1 μL nuclease free water, 0.45 μL forward and reverse primers (concentration 10 pmol/μL) and 1.5 μL DNA template. Cycling conditions were as follows: initial denaturation at 95°C for 5 min, followed by 35 cycles of denaturation (94°C, 30 sec), annealing (55°C, 30 sec), extension (72°C, 90 sec), and a final extension at 72°C for 7 min. PCR products were separated on 1% agarose gels stained with Midori Green Nucleic Acid Staining Solution before UV transillumination. Mapping of results to geographic location was undertaken using QGIS v3.28.2 (53).

## 5. Results

### General features of Erwimp genome

During an attempt to sequence the complete genome of a common Rickettsia endosymbiont in *Culicoides impunctatus* (20), a circular contig with homology to Erwiniaceae bacteria was also identified. The assembled chromosome of Erwimp is 3,461,605 bp in size with an average GC content of 48.2% and an average depth of coverage of 459×, comparable to a 296x coverage for the detected Rickettsia endosymbiont (Supplementary File S3). A BUSCO completeness score of 99.7% (Single-copy: 99.7%, Duplicated: 0%, Fragmented: 0.3%, Missing: 0%) suggests a high-quality genome (Supplementary File S4). Genome annotation found 3,418 protein coding sequences (CDSs) with an average length of 864 bp, 7 copies each of the 5S, 16S and 23s rRNA genes and 69 tRNA genes with a coding density of 85.3%. Of the 3,418 predicted CDSs, 2472 (∼72%) were annotated with putative functions, while 946 (∼28%) CDSs were annotated as hypothetical proteins. In addition, one 132kbp plasmid (pErwimp001) was also identified.

Through an initial phylogenetic screen of 5 concatenated house-keeping genes, the bacterial chromosome appeared to cluster within the *Erwinia* genus of the family (Supplementary File S2). Subsequently, to confirm this as a new species member of the group, the phylogenomic relationship of Erwimp relative to other Erwiniaceae was estimated (Figure 1A) from a set of 144 single-copy orthologues identified among 44 draft or complete genomes (23 *Erwinia*, 15 *Pantoea*, 3 *Izhakiella*, 2 *Tatumella* and 1 *Mixta*). The maximum-likelihood tree placed Erwimp within a subclade of the genus incorporating *Erwinia tracheiphila*, Erwinia psidii, Erwinia mallotivora and Erwinia endophytica, with the most recent known ancestor being Erwinia sp.1945, an endohyphal bacteria of *Microdiplodia* sp.. As an adjunct to the phylogenomic data, Average Nucleotide Identity (ANI) was compared between all genomes. The closest percentage identity of any genome to Erwimp was 74% clearly delineating it as a new species (Figure 1D).

**Figure 1.**
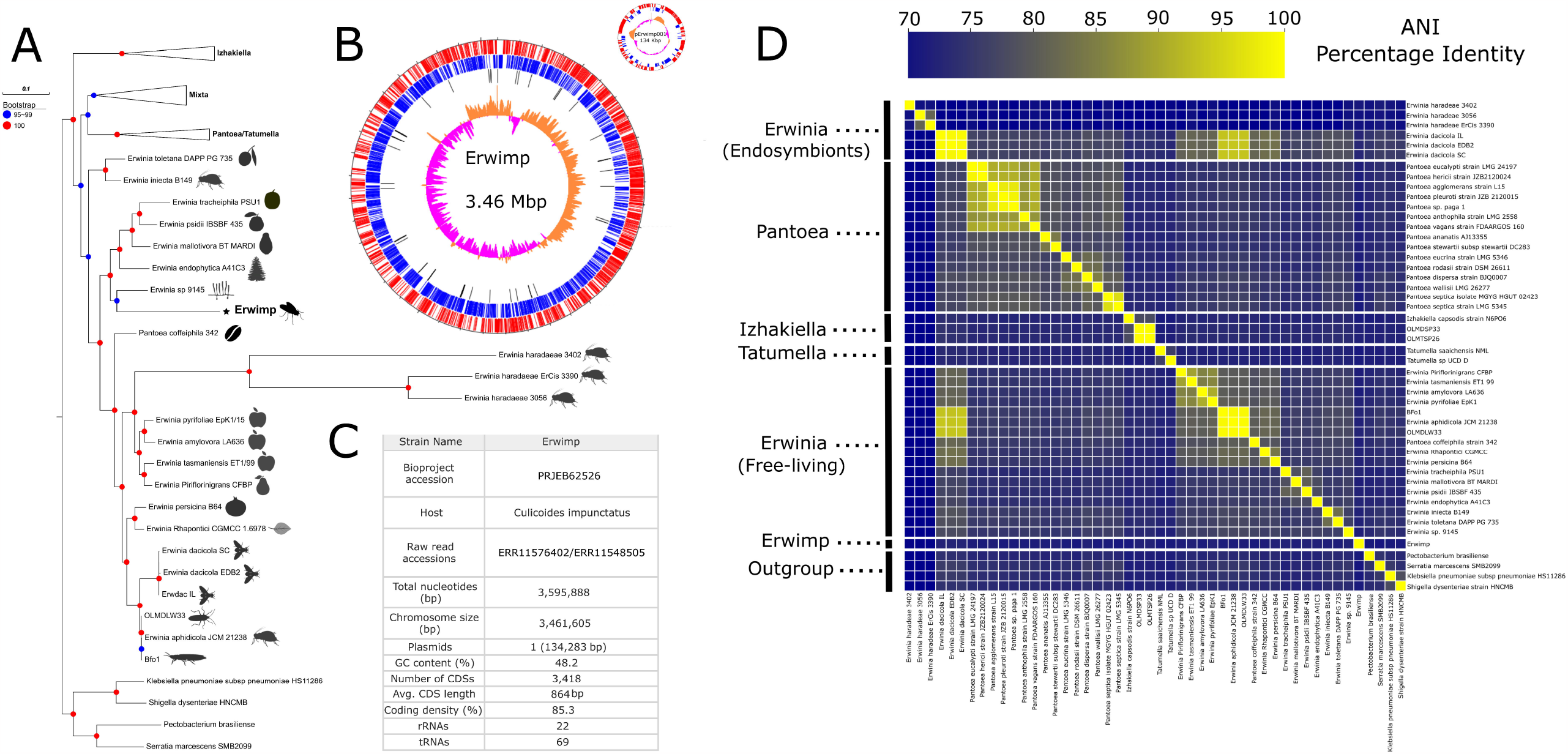
**A)** Maximum-likelihood phylogenomic tree of the Erwiniaceae family based on 144 concatenated single-copy orthologues. Leaf silhouettes represent the isolation source of *Erwinia* spp. strains. The Erwimp isolate described in this study is denoted in bold and a star symbol. **B)** A genome plot of Erwimp. From outermost rings inward, CDSs on the direct strand, CDSs on the reverse strand, tRNAs, G-C skew.**C)** Genome features and project accession information **D)** Average Nucleotide Identity (ANI) heatmap of Erwiniaceae genomes.

### COG, metabolic and pseudogene profiles of Erwimp

Insect endosymbionts and free-living bacteria tend to have different genome compositions as a result of varying evolutionary dynamics (e.g., Muller’s ratchet). Subsequently, to provide insights into the lifestyle of Erwimp, its Clusters of Orthologous Genes (COG) profile was compared to other *Erwinia* (Figure 2A). Generally, Erwimp matched the profiles of both endophytic and epiphytic species with relative enrichment compared to insect endosymbionts of amino acid metabolism and transport (COG category E), signal transduction mechanisms (T), inorganic ion transport (P), and carbohydrate metabolism and transport (G). For both *Erwinia dacicola* and *Erwinia haradaeae* endosymbionts, certain COG categories were enriched in comparison to Erwimp and the free-living *Erwinia*. These included, for *Erwinia haradaeae*, increased proportions of energy production and conversion (C), coenzyme C metabolism and transport (H), translation (J), replication and repair (L), cell wall biogenesis (M) and post-translational modifications (O). Whereas, for *Erwinia dacicola*, increased relative proportions were seen of replication and repair (L) and mobilome genes (X).

**Figure 2.**
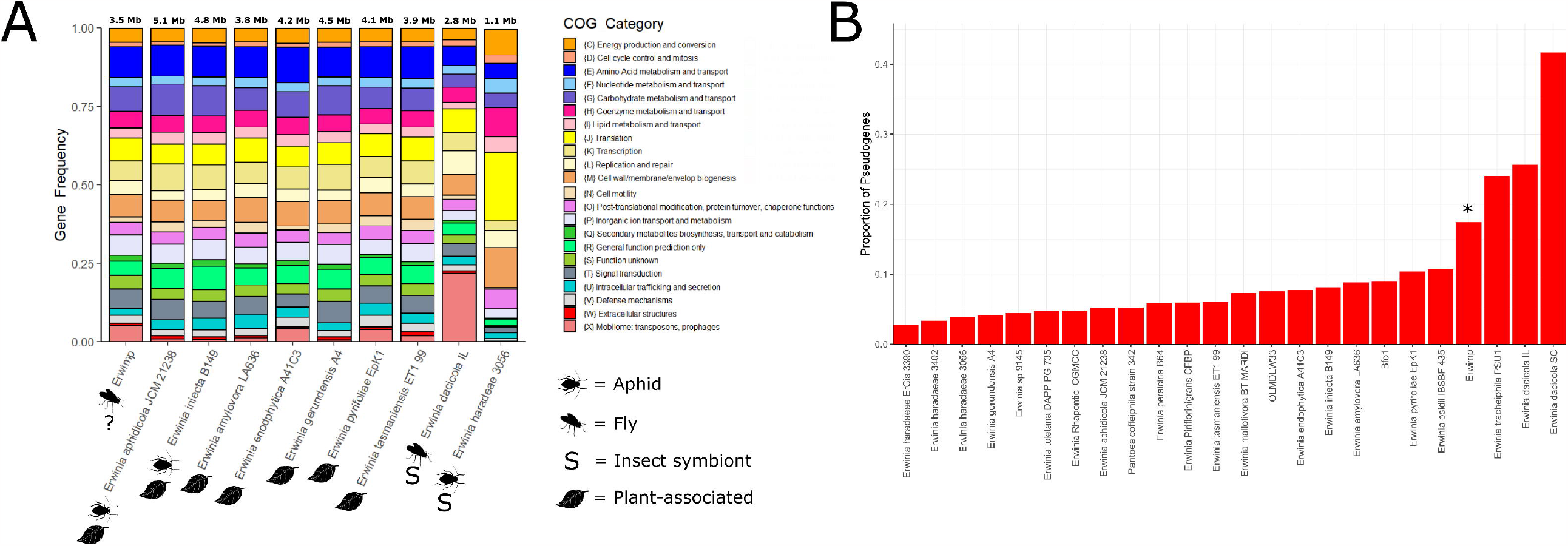
**A)** Frequencies of Clusters of Orthologous Genes (COG) clusters for Erwimp compared to a subset of free-living *Erwinia* strains and insect endosymbionts. **B)** The proportion of pseudogenes attributed to each *Erwinia* species.

Erwimp’s metabolic potential (Figure 3) is somewhat similar to members of the genus which have undergone genome decay as a result of host specialisation (e.g., *Erwinia tracheiphila, Erwinia dacicola* and *Erwinia haradaeae*). For example, amino acid biosynthesis pathways for glutamate and betaine are absent for Erwimp, *Erwinia tracheiphila, Erwinia dacicola* and *Erwinia haradaeae*, but are generally complete elsewhere (Figure 3A). The same pattern is observed for lipid metabolism with absent pathways for beta-oxidation in acyl-coA degradation as well as ketone body synthesis (Figure 3D). Erwimp also lacks the capability of producing UDP-galactose, a key component of nucleotide sugar metabolism (Figure 3C). Co-factors and vitamin pathways were also assessed (Figure 3B) due to both plant and blood-sucking insect bacteria assisting in the provisioning of B-vitamins often lacking in the host’s diet. Erwimp contains the complete pathways for pantothenate, biotin and riboflavin biosynthesis although this does not appear to be unique across the genus, with all free-living species also containing these pathways, but not endosymbionts (apart from *Erwinia haradaeae* and riboflavin). Further evidence of relative genome degradation comes from the proportion of pseudogenes in the Erwimp genome (17.4%), which is higher than most free-living (mean=7.8%) species although slightly lower than those thought to have undergone host-specialisation (e.g., *Erwinia tracheiphila* [24%]; Figure 2B).

**Figure 3.**
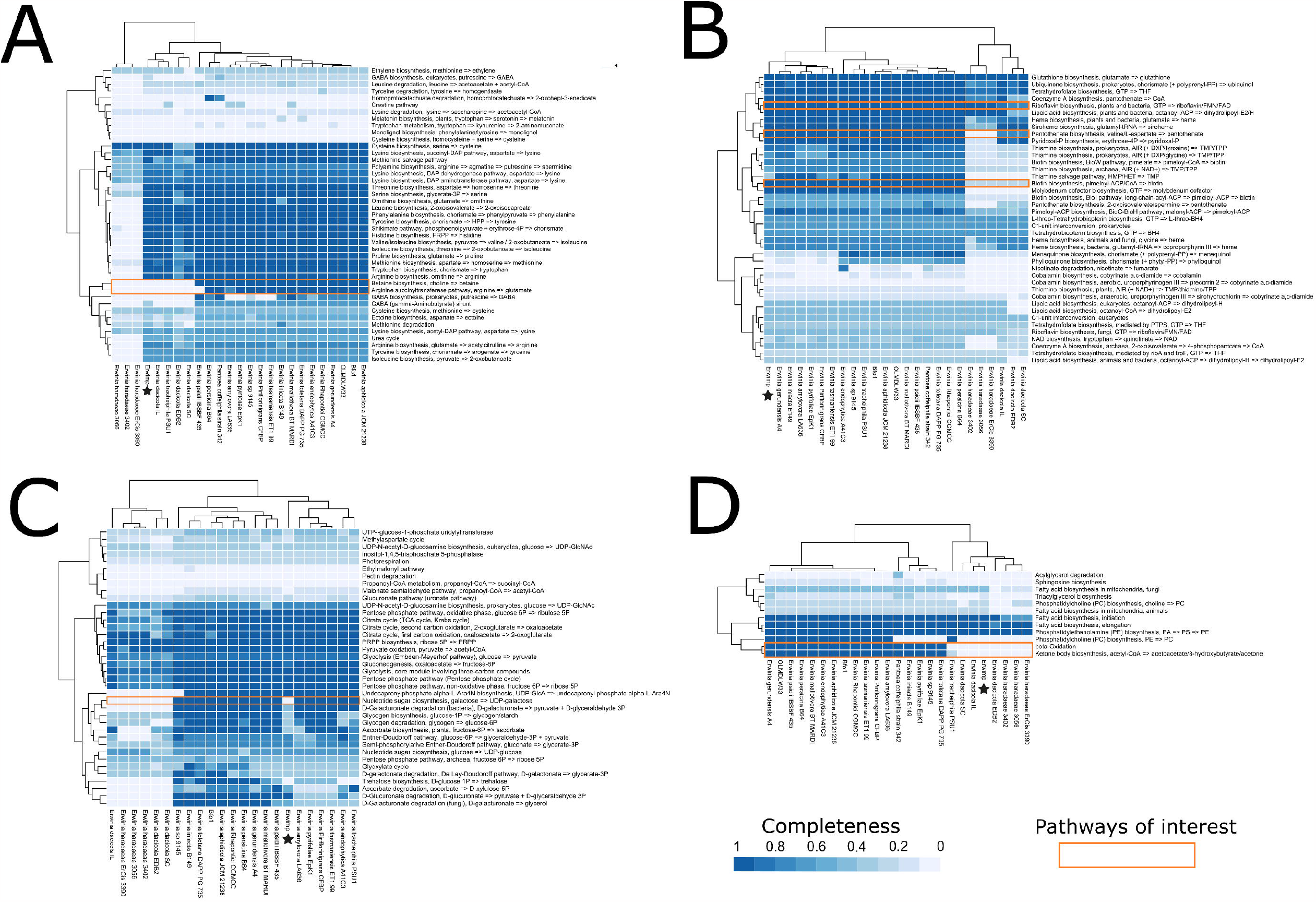
Metabolic profile heatmaps of predicted Kyoto Encyclopedia of Genes and Genomes (KEGG) pathways. Erwimp metabolic profiles are denoted by stars. **A)** Amino acid metabolism. **B)** Vitamins and co-factors. **C)** Carbohydrate metabolism. **D)** Lipid metabolism.

### Genetic repertoire of Erwimp

To identify further evidence that Erwimp has a free-living component to its lifestyle, common *Erwinia* genes relating to virulence and adherence were searched for (Figure 4A). Erwimp appears to hold the capacity to communicate through quorum sensing (QS), containing the QS regulatory system *EsaI/EsaR*, as well as s-ribosylhomocysteine lyase (LuxS) which synthesises autoinducer-2. Additionally, the genome contains a full Type 1 fimbrial gene cassette which is not present in any other members of the genus assessed. Out of the characterised *Erwinia* biofilm exopolysaccharides, amylovoran and levan, the latter but not former is predicted to be produced by Erwimp. Furthermore, a Type VI secretion system spanning across two loci is present although other secretion systems commonly found across the genus (Type II and III) are incomplete or absent. Flagellar genes encoding the filament, rod, hook and P/L ring structures are present, but only *FliMN* genes of the motor apparatus exist (missing basal MS and C ring components) suggesting an absent or imperfect functioning motility system. Intriguingly, a unique Erwimp gene identified is a chitooligosaccharide deacetylase *(ChbG)* homolog. *ChbG* hydrolyzes the N-acetyl group at the reducing-end of chitin disaccharides and is complemented by the presence of a putative chitoporin transport channel *(ChiP)*. Finally, a full complement of lipopolysaccharide (LPS) genes (Lipid A, core and O-antigen) are present in most free-living *Erwinia*, as well as Erwimp.

**Figure 4.**
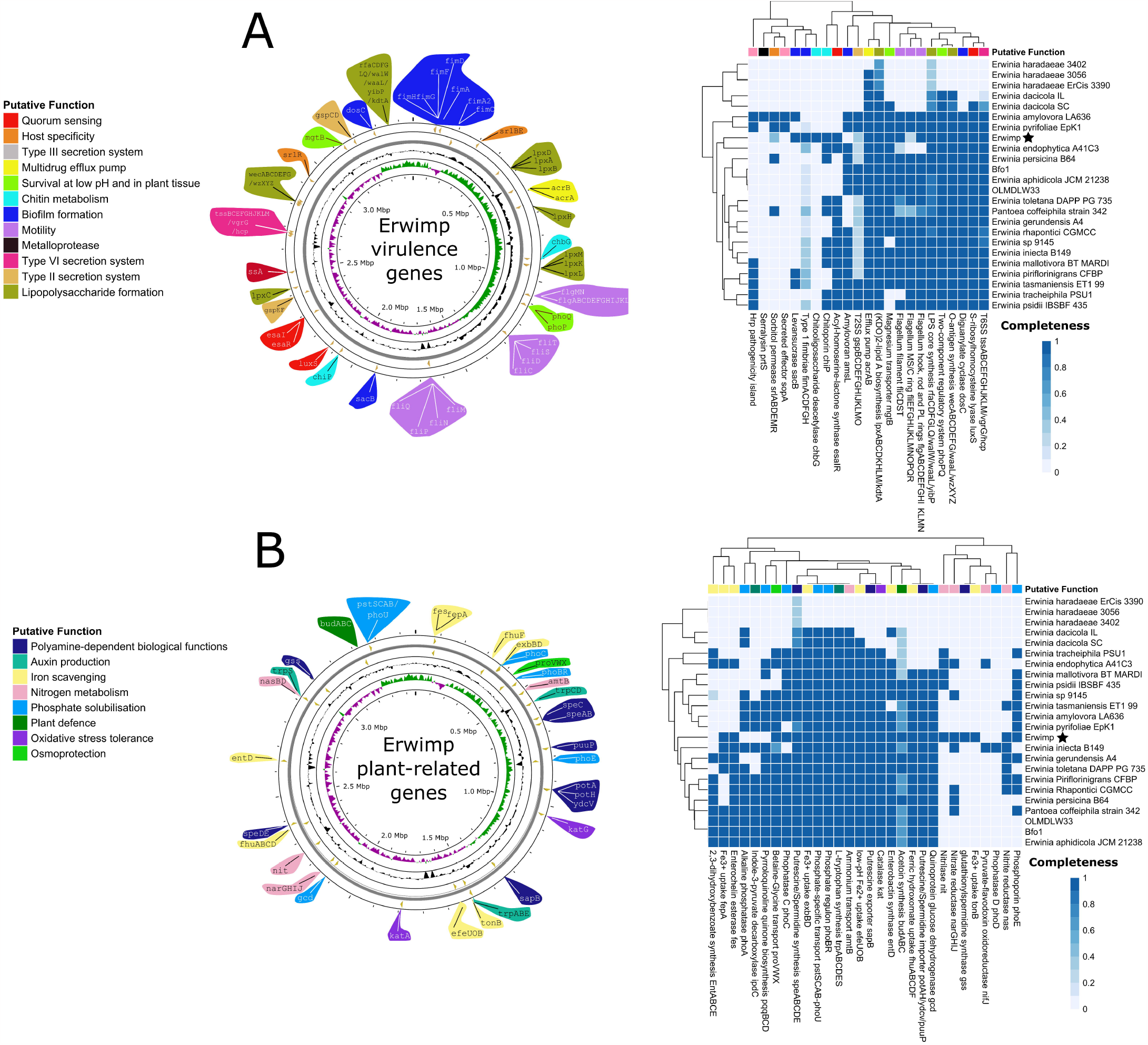
Annotated Erwimp chromosomes detailing common *Erwinia* virulence factors **(A)** and putative plant-related genes **(B)** alongside their completeness in other *Erwinia* species.

Plant-related genes were also assessed to clarify potential plant-growth and protection attributes (Figure 4B). Unlike *Erwinia* endosymbionts, Erwimp contains all necessary genes to produce *(SpeABCDE)* and import/export spermidine *(PuuP, SapB, PotA, PotH, YdcV)* suggesting a role in polyamine-related processes such as swarming in bacteria and potentially plant defence responses and abiotic stress tolerance. Additionally, osmoprotectant (betaine glycine transporter; *ProVWX)* and antioxidant (Catalase; *Kat*) genes present exclusively in free-living plant *Erwinia* are also found in Erwimp. Furthermore, a biosynthetic gene cluster for the plant-protecting volatile organic compound acetoin *(BudABC)* is present.

Regarding the capacity of Erwimp to mobilise nutrients for potential plant hosts, Erwimp appears to have lost key genes relating to phosphate solubilisation present in other plant-associated *Erwinia*. These include genes producing pyrroloquinoline quinone *(PqqBCD)*, a cofactor for quinoprotein glucose dehydrogenase *(Gcd)*, enabling the solubilisation of inorganic phosphorous. Partial pathways related to other plant physiological processes are found such as the complete split operon for the biosynthesis of L-tryptophan *(TrpABCDES)*, a precursor of the phytohormone indole-3-acetic acid (IAA). However, the ipdC pathway converting L-tryptophan to IAA is absent in Erwimp. Iron scavenging genes are also found in the genome. Specifically, the chromosome contains the enterobactin synthase gene, *EntD*, enabling the final step in enterobactin synthesis, but lacks the ability to produce the upstream precursor 2,3-dihydroxybenzoic acid. A desferrioxamine transport system *(FhuABCDF)*, responsible for the sequestering of ferric hydroxamate siderophores is also present. With respect to nitrogen metabolism, Erwimp nitrate/nitrite reductases *(NarGHI, NasBD)* and a nitrilase *(Nit)* suggest potential roles in plant nitrogen assimilation but the bacterium lacks nitrogen fixing pathways.

### Carotenoid and Intimin-like genes

Secondary metabolite analysis identified the presence of a carotenoid biosynthetic gene cluster (Figure 5). Only three other *Erwinia* genomes assessed for secondary metabolites were predicted to contain similar operons (Supplementary File S5). Of interest, is the loss of zeaxanthin glucosyltransferase *(CrtX)* within Erwimp’s gene cluster compared to its nearest known relative *Erwinia* sp. 9145. *CrtX* is necessary for the synthesis of zeaxanthin glucosides and suggests Erwimp is only able to synthesise the simpler linear zeaxanthin molecule.

**Figure 5.**
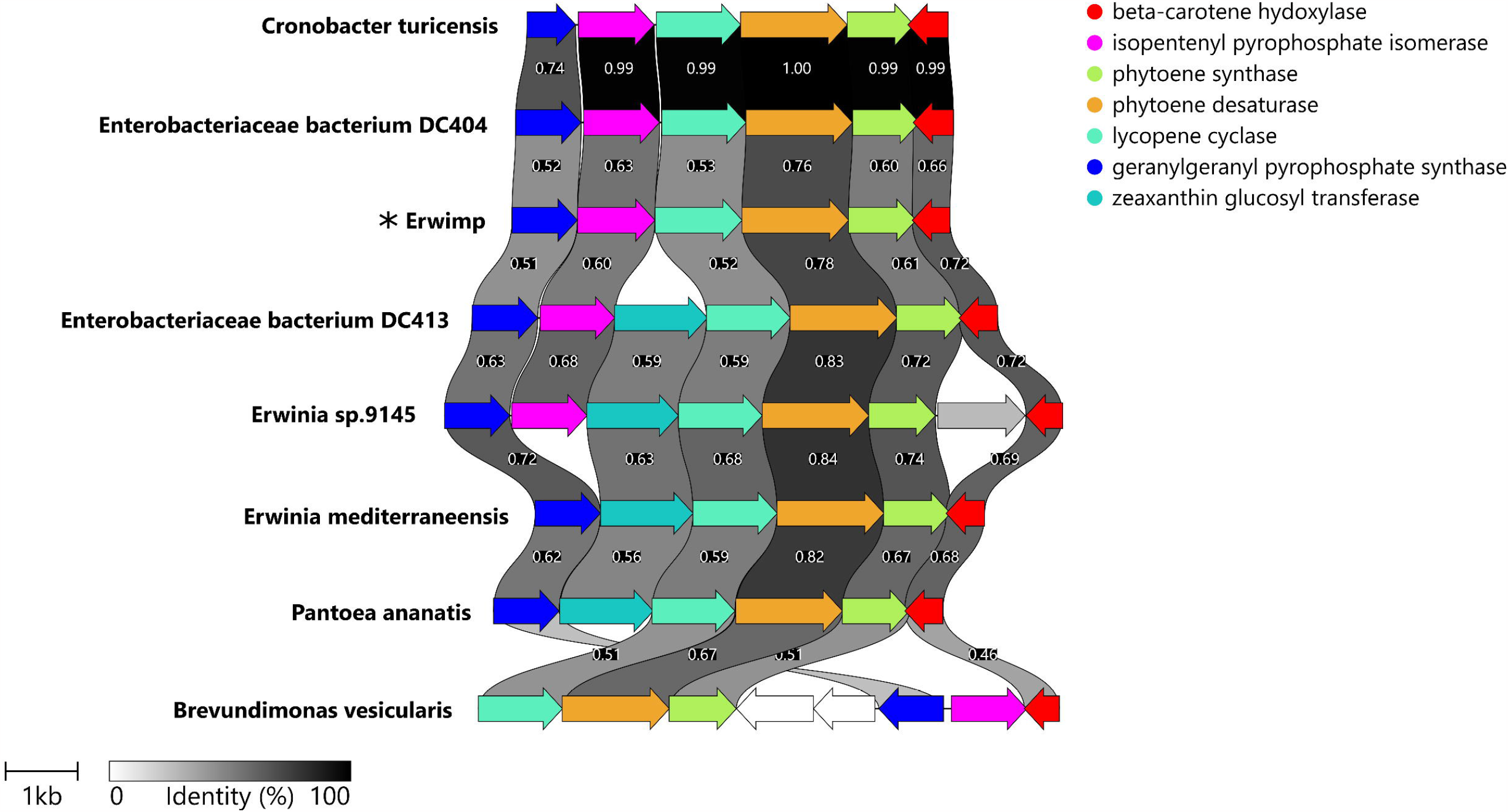
Clinker plot used to compare and visualise the similarity of the Erwimp carotenoid biosynthesis operon uncovered by antiSMASH with other gene clusters based on an all-vs-all similarity matrix.

A 132kbp plasmid (pErwimp001) was also assembled and annotated (Supplementary File S6). pErwimp001 has copies of notable genes present on the chromosome including a putrescine importer *(PuuP)*, an acyl-homoserine lactone synthase/repressor (*EsaI/EsaR*) and fimbriae genes *(FimC, FimD)*. In addition, an ectoine dioxygenase *(EctD)*, catalysing the production of the desiccant protectant hydroxyectoine is present on pErwimp001 but absent in all other *Erwinia* genomes analysed, with the nearest hit being to Izhakiella capsodis (WP_092877528.1; 77% identity). Intriguingly, the largest gene on the plasmid (eae; 3213 amino acid residues) is an intimin homolog. Three copies of this gene are present on the Erwimp chromosome with a varying number of bacterial immunoglobulin-like domains (BIGs) repeats at the C-terminus (Figure 6A). Two of these (Erwimp_eae_1 and Erwimp_eae_3) contain all 3 intimin domains (signal peptide, transmembrane beta barrel and passenger domains) as an operon, whereas Erwimp_eae_2 contains two stop codons between beta and passenger domains suggesting pseudogenization. No other *Erwinia* genomes in the comparative analysis are predicted to contain intimins and the close identity of pErwimp001_eae_1, Erwimp_eae_2 and Erwimp_eae_3 (>97% identity over 2664 amino acid residues) suggests a recent plasmid integration involving the Erwimp chromosome followed by gene duplication. A phylogeny of the conserved beta domain suggests Erwimp_eae_1 has a separate evolutionary origin to the other 3 intimins (Figure 6B).

**Figure 6.**
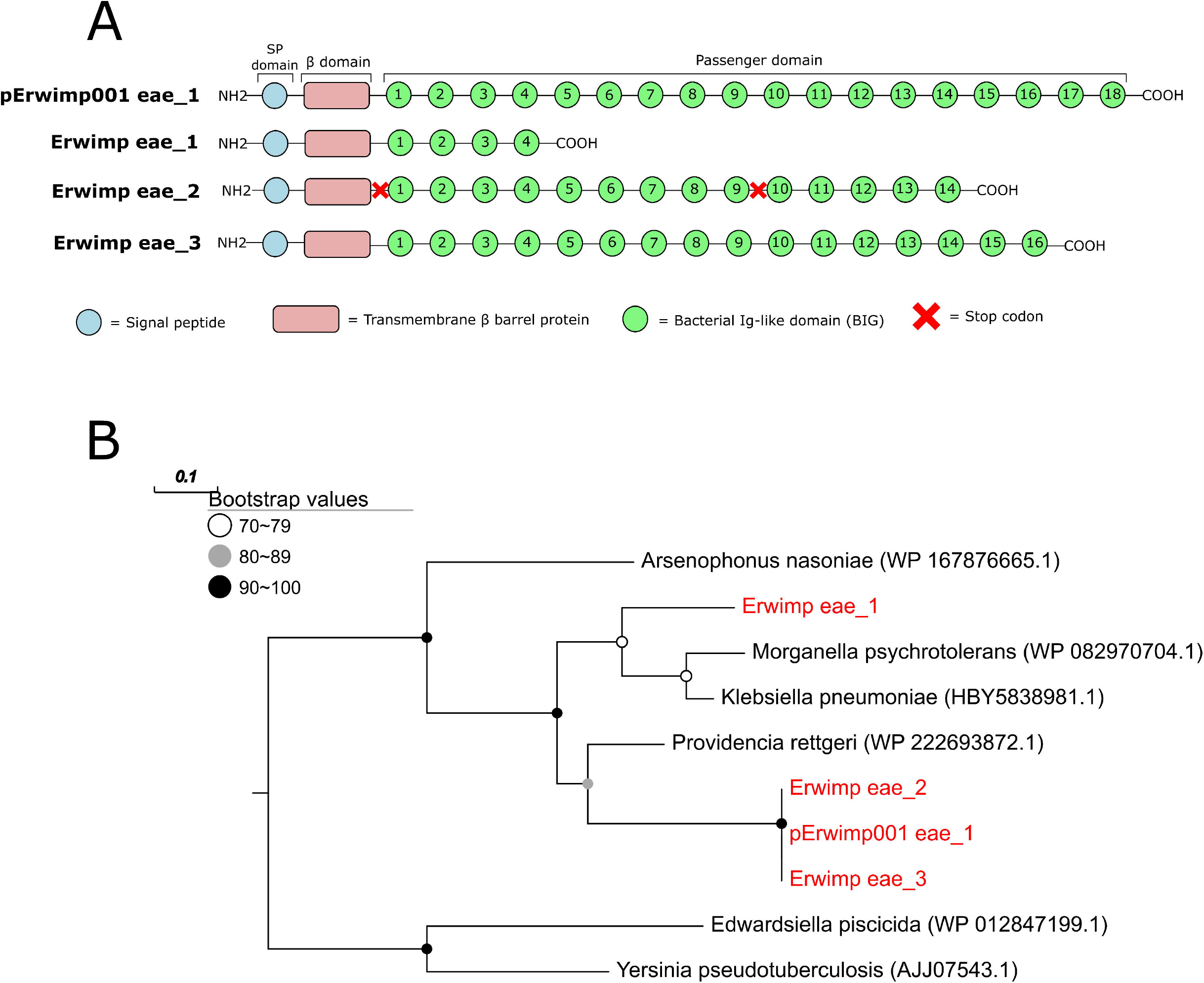
**A)** Interpro predictions of domains found in multiple Intimin-like proteins found on an Erwimp plasmid (pErwimp001_eae_1) and chromosome (Erwimp_eae_1, Erwimp_eae_1, Erwimp_eae_1). **B)** Maximum-likelihood phylogeny of the beta-domain from intimin copies based on a 240 amino acid alignment.

### Targeted screening of *Culicoides impunctatus* individuals

A total of 115 *Culicoides impunctatus* individuals were screened for the presence of *Erwinia*. Of the remaining 51 insects from the 2020 catch, 28 (55%) were positive by conventional PCR. Returning the following year, no infected insects were identified at the same site as well as neighbouring locations of the 64 screened (Figure 7).

**Figure 7.**
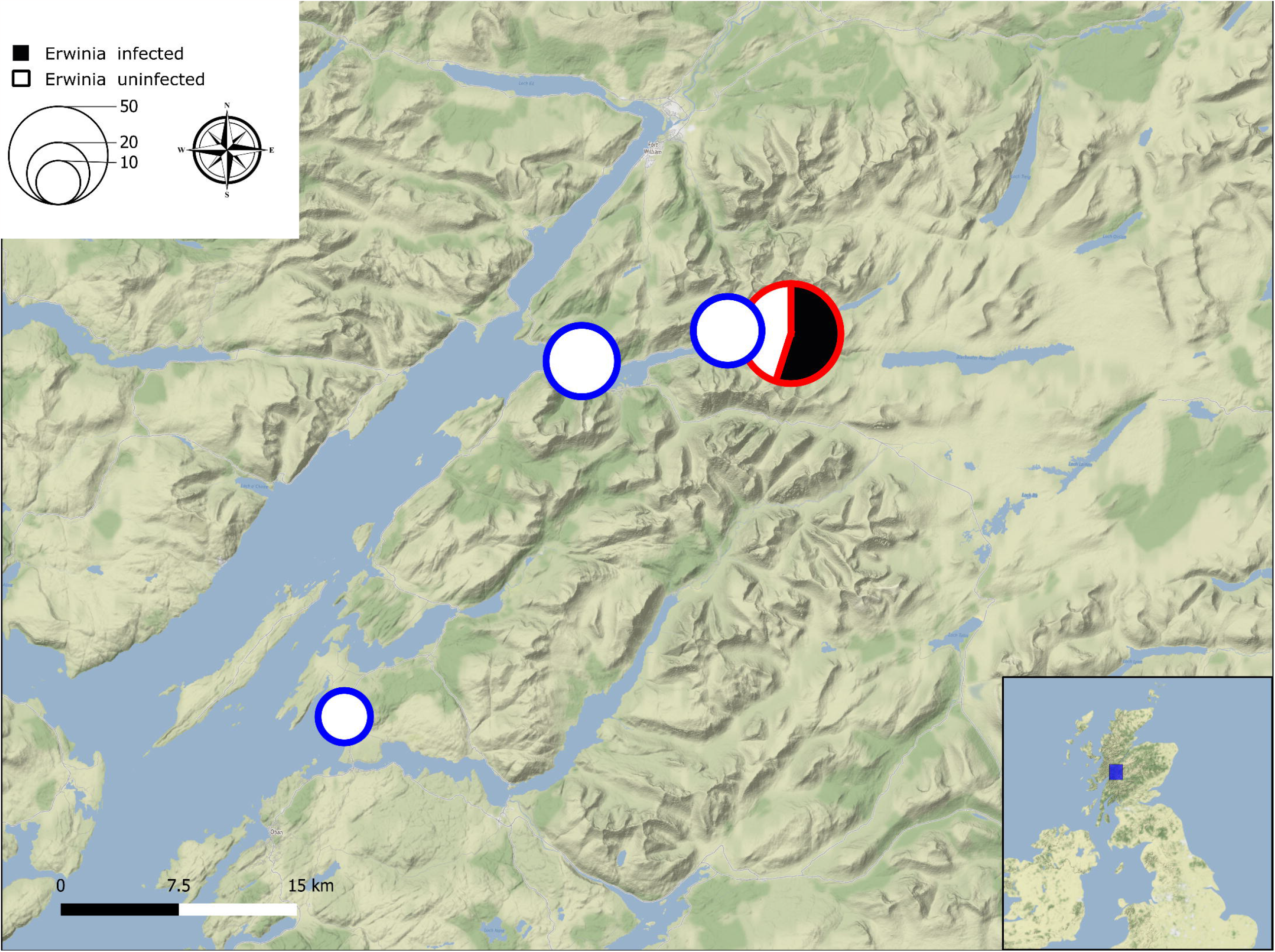
A QGIS map of the Western Scottish highlands depicting catch sites for *Culicoides impunctatus* during field catches between September 2020 (red) and July 2021 (blue) for the targeted PCR screening of *Erwinia*. Size of circles indicate number of specimens per site with the black portion of a circle showing the number of PCR positives.

### Metagenome assessment of potential Erwimp hosts

Aside from *Culicoides impunctatus*, SSU rRNAs exploration of the BGI-derived metagenomic raw-read data provided evidence for one potential plant host, Arabidopsis thaliana (Brassicaceae) (Supplementary File S7). Additional eukaryotes found in the dataset include the fungi *Entomophthora culicis* (Entomophthoraceae) and *Kluyveromyces lactis* (Saccharomycetaceae).

## 6. Discussion

Interactions between microbes and their insect hosts can be transitory or sustained depending on their functional relationship (1,54). In the case of *Erwinia*, members of the genus have evolved to fill various ecological niches including as insect symbionts and phytopathogens (10,15,55). Thus, the discovery of Erwimp in *Culicoides impunctatus* raises questions as to the nature of this association.

The genome of Erwimp is generally comparable to endophytic and epiphytic members of the genus. The similarities in COG profiles (Figure 2A) between Erwimp and other free-living *Erwinia* include the relative enrichment of carbohydrate and amino acid metabolism compared to unculturable insect endosymbionts, and is suggestive of Erwimp’s lack of dependence on a host for basic metabolic processes. A further proxy of lifestyle is the relative number of genomic mobile genetic elements (MGEs). Newton and Bordenstein (56) demonstrated that the largest numbers of MGEs are found in facultative intracellular bacteria (e.g., *Erwinia dacicola*), followed by obligate extracellular (e.g., *Erwinia pyrifoliae*) and obligate intracellular bacteria (e.g., *Erwinia haradaeae*). This pattern is observed in the current study with the mobilome profiles of Erwimp (5% proportion mobilome genes) comparable to *Erwinia* pyrifoliae (4%) and *Erwinia* endophytica (4%) but being much lower than *Erwinia dacicola* (21%) and higher than *Erwinia haradaeae* (0.1%), giving more credence to an obligate extracellular lifestyle for Erwimp. Furthermore, virulence factors specific to free-living *Erwinia* tend to be found in Erwimp, many of which are likely directly or indirectly related to biofilm formation. These include the ability to produce the exopolysaccharide (EPS) levan (57), a type VI secretion system associated with EPS production and interbacterial competition (58), a lipopolysaccharide (LPS), quorum sensing genes and a full type 1 fimbrial gene cluster.

The frequent transmission of *Erwinia* to plant hosts via insect vectors (13,14) suggests local vegetation should be considered as additional putative hosts of Erwimp. Potential plant hosts include *Arabidopsis thaliana* which occurs predominantly in the Scottish highlands during the summer months (59) at the same time as midge season and was found as the only plant SSU RNA detected in the metagenomic data (Supplementary File S7). Further hosts to consider include those part of *C. impunctatus*’ larval/pupal habitats, such as *Sphagnum* spp. and Polytrichum spp. mosses (60), as well as adult resting sites like downy birch (*Betula pubescens*) trees (61). Homologs of putative plant growth promoting and defence genes were observed in the Erwimp genome (Figure 4B), however, these genes often included those with an ambiguous ontology. For example, spermidine/putrescine synthesis and transport genes (*speABCDE/puuP, sapB, potA, potH, ydcV*) could be related to host or bacterial polyamine-related functions such as abiotic stress tolerance (62) in the former and chemotaxis or growth in the latter (63). Additionally, genes related to nitrogen assimilation, siderophore synthesis and phosphate mobilization are present but Erwimp still lacks full pathways for enterobactin synthesis, nitrogen fixation, IAA synthesis and phosphate solubilization by gluconic acid.

Despite this, a notable carotenoid biosynthetic gene cluster, implicated as a virulence factor in plant pathogens of the sister genus, Panotea (64–66), is also present in Erwimp (Figure 5). Carotenoids are pigments associated with UV and oxidative stress tolerance in Pantoea spp. where they are under regulation by the *EsaI/EsaR* quorum-sensing regulatory system (64). Aside from assisting in photo tolerance, phytoene synthase (crtB) defective mutants have been shown to impair biofilm formation (65) and root colonization (66) in Pantoea sp. YR343. Furthermore, the predicted product of this operon, zeaxanthin, is a precursor to the phytohormone abscisic acid (ABA) (67). Thus, despite Erwimp’s loss of the idpC pathway for IAA synthesis, assistance in ABA production may still play a role in the growth control of a plant host.

Although *Erwinia* appear to be primarily associated with plants and insects, there are examples of endohyphal *Erwinia* (16); these include *Erwinia* sp.1945, found in Microdiplodia sp. and the closest known relative of Erwimp. Other putative hosts detected in the metagenome include the fungal insect pathogen, *Entomophthora culicis* and the yeast *Kluyveromyces lactis*. Of particular intrigue is the presence of a chitooligosaccharide deacetylase *(ChbG)* with a possible role either in a fungal or insect niche. ChbG is involved in the production of N-acetylglucosamine (GlcNAc-GlcN) from the chitin disaccahride N,N’-diacetylchitobiose, to assist in its import across the bacterial periplasm (68). The presence of a chitoporin also indicates the utilisation of chitin as an energy source possibly from fungal cell walls or the peritrophic membrane of insect midguts. In addition, GlcNAc-GlcN acts as a chemoattractant for *Vibrio* (69) suggesting *ChbG* may have an additional role in swarming behaviour.

Regardless of additional Erwimp hosts, the genome’s large proportion of pseudogenes (Figure 2B) along with the loss of certain components of amino acid, carbohydrate, and fatty acid metabolism (Figure 3) is reminiscent of *Erwinia tracheiphila*. In this case, the cucurbit plant wilt pathogen, has become obligately insect-transmitted leading to genome decay (70) and suggests a possible similar host restriction for Erwimp. Intriguingly, multiple intimin (eae) copies unique to *Erwinia* are found within the Erwimp genome (Figure 6). Intimins are a family of outer membrane proteins generally found in the Enterobacterales which act as adhesins (71). The presence of three near identical intimin-like proteins on the plasmid and chromosome (2 intact and one pseudogenised), indicates a recent chromosome integration event, and suggests these are important for host interactions. Aside from offering a further putative mechanism for Erwimp adherence to eukaryotic cells and biofilm formation (72), in vivo and ex vivo studies investigating the interchange of *E. coli* intimin subtypes (73), found these adherence factors are determinants of host specificity and tissue tropisms offering further evidence for a narrow host range for Erwimp.

Of the culturable *Erwinia* strains associated with insects, several inhabit the gut and have been shown to be sustained over several generations (e.g., BFo1 symbiont of western flower thrips [74]) or are transient associations (e.g., *Erwinia* aphidicola and *Erwinia* iniecta in aphids [75,76]). After the initial identification of Erwimp at intermediate prevalence in the 2020 collection, the acquisition of more infected *C. impunctatus* for biochemical and imaging analysis was attempted the following field season (2021), however, no more Erwimp positive insects were identified (Figure 7). This suggests Erwimp may inhabit *C. impunctatus* as a transient host, as co-evolved symbioses between gut bacteria and insects are often widespread and at high prevalence in populations (77).

*Culicoides impunctatus* impacts Scottish tourism and forestry industries through its voracious biting of humans and as a hazard to safe arboricultural operations (78). Given the abundance and importance of *C. impunctatus* as a biting pest, the effects of this bacterium on insect fitness is of great interest. Prokaryotes have been shown to supplement their insect hosts with B-vitamins, particularly those reliant on blood or phloem feeding (6). Although Erwimp contains the full pathways for the synthesis of several B-vitamins (Figure 3.), this is not an exclusive feature amongst other strains indicating this is likely not a symbiotic feature. Experimental assessment of host fitness for several *Erwinia* present in insect guts have provided a spectrum of outcomes from pathogenic to mutualistic (76, 79, 80). However, of the members deemed as pathogens (*Erwinia* iniecta, *Erwinia* aphidicola) experimentation was conducted under laboratory conditions and possibly at unrealistic titres using an artificial diet (76,79). Similarly, the beneficial effects on development and fecundity observed with BFo1 in western flower thrips is diet-dependent (80). Overall, the cryptic nature of *Erwinia*-insect interactions suggest further work is required to assess the impact of Erwimp on *Culicoides impunctatus*, beginning with confirming culturability. Furthermore, consideration should be given to the insect vector potential of *C. impunctatus* to transmit Erwimp, as this is likely to have an impact on local vegetation with both plant pathogenicity and growth promotion (81) possible.

## 7 Author statements

### 7.1 Author contributions

JP is the sole author of this work and was responsible for the experimental design, experiments, data analysis, and writing/interpretation of the data.

### 7.4 Ethical approval

Not Applicable

## 7.5 Acknowledgements

The author would like to thank Prof. Greg Hurst for giving critical feedback for this preprint.

## 7.6 Data accessibility

Assemblies and raw sequencing data can be found under Bioproject PRJEB62526 in the European Nucleotide Archive. Bioinformatic and figure code can be found in Supplementary File S3.

## References

1. Ferrari J, Vavre F. Bacterial symbionts in insects or the story of communities affecting communities. Phil Trans R Soc B. 2011;366(1569):1389–400.

2. Engelstädter J, Hurst GDD. The ecology and evolution of microbes that manipulate host reproduction. Annu Rev Ecol Evol Syst. 2009;40(1):127–49.

3. Teixeira L, Ferreira Á, Ashburner M. The bacterial symbiont Wolbachia induces resistance to RNA viral infections in Drosophila melanogaster. PLoS Biol. 2008;6(12):e1000002.

4. Hamilton PT, Peng F, Boulanger MJ, Perlman SJ. A ribosome-inactivating protein in a Drosophila defensive symbiont. Proc Natl Acad Sci U S A. 2016;113(2):350–5.

5. Douglas AE. Nutritional Interactions in Insect-Microbial Symbioses: Aphids and their symbiotic bacteria Buchnera. Annu. Rev. Entomol. 1998;43(1):17–37.

6. Manzano-Marín A, Oceguera-Figueroa A, Latorre A, Jiménez-García LF, Moya A. Solving a bloody mess: B-vitamin independent metabolic convergence among Gammaproteobacterial obligate endosymbionts from blood-feeding arthropods and the leech Haementeria officinalis. Genome Biol Evol. 2015;7(10):2871–84.

7. Sela S, Nestel D, Pinto R, Nemny-Lavy E, Bar-Joseph M. Mediterranean fruit fly as a potential vector of bacterial pathogens. Appl Environ Microbiol. 2005;71(7):4052–6.

8. Picciotti U, Araujo Dalbon V, Ciancio A, Colagiero M, Cozzi G, De Bellis L, et al. “Ectomosphere”: Insects and microorganism interactions. Microorganisms. 2023;11(2):440.

9. Menelas B, Block CC, Esker PD, Nutter FW. Quantifying the feeding periods required by Corn Flea Beetles to acquire and transmit Pantoea stewartii. Plant Disease. 2006;90(3):319–24.

10. Manzano-Marín A, Coeur D’Acier A, Clamens AL, Orvain C, Cruaud C, Barbe V, et al. Serial horizontal transfer of vitamin-biosynthetic genes enables the establishment of new nutritional symbionts in aphids’ di-symbiotic systems. ISME J. 2020;14(1):259–73.

11. Estes AM, Hearn DJ, Bronstein JL, Pierson EA. The Olive Fly Endosymbiont, “Candidatus *Erwinia dacicola*,” switches from an intracellular existence to an extracellular Existence during host insect development. Appl Environ Microbiol. 2009;75(22):7097–106.

12. Ben-Yosef M, Pasternak Z, Jurkevitch E, Yuval B. Symbiotic bacteria enable olive flies (Bactrocera oleae ) to exploit intractable sources of nitrogen. J Evol Biol. 2014;27(12):2695–705.

13. Emmett BJ, Baker LAE. Insect transmission of Fireblight. Plant Pathology. 1971;20(1):41–5. 14.

14. Sasu MA, Seidl-Adams I, Wall K, Winsor JA, Stephenson AG. Floral transmission of *Erwinia tracheiphila* by Cucumber Beetles in a wild Cucurbita pepo. Environ Entomol. 2010 Feb 1;39(1):140–8.

15. Palacio-Bielsa A, Roselló M, Llop P, López MM. *Erwinia* spp. from pome fruit trees: similarities and differences among pathogenic and non-pathogenic species. Trees. 2012 Feb;26(1):13–29.

16. Baltrus DA, Dougherty K, Arendt KR, Huntemann M, Clum A, Pillay M, et al. Absence of genome reduction in diverse, facultative endohyphal bacteria. Microb Genom. 2017;3(2):e000101.

17. O’Hara CM, Steigerwalt AG, Hill BC, Miller JM, Brenner DJ. First report of a human isolate of *Erwinia* persicinus. J Clin Microbiol. 1998;36(1):248–50.

18. Shin SY, Lee MY, Song JH, Ko KS. New *Erwinia*-like organism causing cervical lymphadenitis. J Clin Microbiol. 2008;46(9):3156–8.

19. Boorman J, Goddard P. Observations on the biology of *Culicoides impunctatus* Goetgh. (Dipt., Ceratopogonidae) in southern England. Bull Entomol Res. 1970;60(2):189–98.

20. Davison HR, Pilgrim J, Wybouw N, Parker J, Pirro S, Hunter-Barnett S, et al. Genomic diversity across the Rickettsia and ‘Candidatus Megaira’ genera and proposal of genus status for the Torix group. Nat Commun. 2022;13(1):2630.

21. De Coster W, D’Hert S, Schultz DT, Cruts M, Van Broeckhoven C. NanoPack: visualizing and processing long-read sequencing data. Bioinformatics. 2018;34(15):2666–9.

22. Kolmogorov M, Yuan J, Lin Y, Pevzner PA. Assembly of long, error-prone reads using repeat graphs. Nat Biotechnol. 2019;37(5):540–6.

23. Chen Y, Chen Y, Shi C, Huang Z, Zhang Y, Li S, et al. SOAPnuke: a MapReduce accelerationsupported software for integrated quality control and preprocessing of high-throughput sequencing data. GigaScience. 2018;7(1):1–6.

24. Li D, Liu CM, Luo R, Sadakane K, Lam TW. MEGAHIT: an ultra-fast single-node solution for large and complex metagenomics assembly via succinct de Bruijn graph. Bioinformatics. 2015;31(10):1674–6.

25. Parks DH, Imelfort M, Skennerton CT, Hugenholtz P, Tyson GW. CheckM: assessing the quality of microbial genomes recovered from isolates, single cells, and metagenomes. Genome Res. 2015;25(7):1043–55.

26. Bushnell B. BBMap: a fast, accurate, splice-aware aligner. 2014. BBMap: a fast, accurate, spliceaware aligner. Available at: https://sourceforge.net/projects/bbmap.

27. Li H, Handsaker B, Wysoker A, Fennell T, Ruan J, Homer N, et al. The Sequence Alignment/Map format and SAMtools. Bioinformatics. 2009;25(16):2078–9.

28. Walker BJ, Abeel T, Shea T, Priest M, Abouelliel A, Sakthikumar S, et al. Pilon: An integrated tool for comprehensive microbial variant detection and genome assembly improvement. PLoS ONE. 2014;9(11):e112963.

29. Manni M, Berkeley MR, Seppey M, Zdobnov EM. BUSCO: Assessing genomic data quality and beyond. Curr Protoc. 2021;1(12):e323.

30. Nishimura O, Hara Y, Kuraku S. gVolante for standardizing completeness assessment of genome and transcriptome assemblies. Bioinformatics. 2017;33(22):3635–7.

31. Seemann T. Prokka: rapid prokaryotic genome annotation. Bioinformatics. 2014;30(14):2068–9. 32.

32. Blin K, Shaw S, Augustijn HE, Reitz ZL, Biermann F, Alanjary M, et al. antiSMASH 7.0: new and improved predictions for detection, regulation, chemical structures and visualisation. Nucleic Acids Res. 2023;51(W1):W46–50.

33. Eren AM, Kiefl E, Shaiber A, Veseli I, Miller SE, Schechter MS, et al. Community-led, integrated, reproducible multi-omics with anvi’o. Nat Microbiol. 2020;6(1):3–6.

34. Castresana J. Selection of conserved blocks from multiple alignments for their use in phylogenetic analysis. Mol Biol and Evol. 2000;17(4):540–52.

35. Kalyaanamoorthy S, Minh BQ, Wong TKF, Von Haeseler A, Jermiin LS. ModelFinder: fast model selection for accurate phylogenetic estimates. Nat Methods. 2017;14(6):587–9.

36. Nguyen LT, Schmidt HA, Von Haeseler A, Minh BQ. IQ-TREE: A Fast and effective stochastic algorithm for estimating maximum-likelihood phylogenies. Mol Biol Evol. 2015;32(1):268–74.

37. Pritchard L, Glover RH, Humphris S, Elphinstone JG, Toth IK. Genomics and taxonomy in diagnostics for food security: soft-rotting enterobacterial plant pathogens. Anal Methods. 2016;8(1):12–24.

38. Wickham H. ggplot2: Elegant graphics for data analysis. 2009. Available from: https://link.springer.com/10.1007/978-0-387-98141

39. R studio team. RStudio: Integrated development for R. 2022. Available from: http://www.rstudio.com/

40. Syberg-Olsen MJ, Garber AI, Keeling PJ, McCutcheon JP, Husnik F. Pseudofinder: Detection of pseudogenes in prokaryotic genomes. Mol Biol Evol. 2022;39(7):msac153.

41. Veseli I, Chen YT, Schechter MS, Vanni C, Fogarty EC, Watson AR, et al. Microbes with higher metabolic independence are enriched in human gut microbiomes under stress. bioRxiv [Preprint] Available from: http://biorxiv.org/lookup/doi/10.1101/2023.05.10.540289

42. Kolde R. pheatmap. 2018. Available from: https://cran.r-project.org/web/packages/pheatmap/

43. Gruber-Vodicka HR, Seah BKB, Pruesse E. phyloFlash: Rapid small-subunit rRNA profiling and targeted assembly from metagenomes. mSystems. 2020;5(5):e00920–20.

44. Piqué N, Miñana-Galbis D, Merino S, Tomás J. Virulence factors of *Erwinia* amylovora: A review. IJMS. 2015;16(12):12836–54.

45. Bruto M, Prigent-Combaret C, Muller D, Moënne-Loccoz Y. Analysis of genes contributing to plant-beneficial functions in plant growth-promoting rhizobacteria and related Proteobacteria. Sci Rep. 2014;4(1):6261.

46. Karp PD, Paley S, Krummenacker M, Kothari A, Wannemuehler MJ, Phillips GJ. Pathway Tools management of pathway/genome data for microbial communities. Front Bioinform. 2022;2:869150.

47. Grant JR, Enns E, Marinier E, Mandal A, Herman EK, Chen C yu, et al. Proksee: in-depth characterization and visualization of bacterial genomes. Nucleic Acids Res. 2023;51(W1):W484–92.

48. Project Inkscape. 2020. Available from: https://inkscape.org

49. Gilchrist CLM, Chooi YH. Clinker & clustermap.js: automatic generation of gene cluster comparison figures. Bioinformatics. 2021;37(16):2473–5.

50. Paysan-Lafosse T, Blum M, Chuguransky S, Grego T, Pinto BL, Salazar GA, et al. InterPro in 2022. Nucleic Acids Res. 2023;51(D1):D418–27.

51. Pilgrim J, Ander M, Garros C, Baylis M, Hurst GDD, Siozios S. Torix group Rickettsia are widespread in Culicoides biting midges (Diptera: Ceratopogonidae), reach high frequency and carry unique genomic features. Environl Microbiol. 2017;19(10):4238–55.

52. Dallas JF, Cruickshank RH, Linton YM, Nolan DV, Patakakis M, Braverman Y, et al. Phylogenetic status and matrilineal structure of the biting midge, Culicoides imicola, in Portugal, Rhodes and Israel. Med Vet Entomol. 2003;17(4):379–87.

53. QGIS Development Team. QGIS Geographic Information System. 2022. Available from: http://qgis.osgeo.org

54. Coolen S, Rogowska-van der Molen M, Welte CU. The secret life of insect-associated microbes and how they shape insect–plant interactions. FEMS Microbiol Ecol. 2022;98(9):fiac083.

55. Capuzzo C, Firrao G, Mazzon L, Squartini A, Girolami V. ‘Candidatus *Erwinia dacicola*’, a coevolved symbiotic bacterium of the olive fly Bactrocera oleae (Gmelin). International Journal of Systematic and Evol Microbiol. 2005;55(4):1641–7.

56. Newton ILG, Bordenstein SR. Correlations between bacterial ecology and mobile DNA. Curr Microbiol. 2011;62(1):198–208.

57. Koczan JM, McGrath MJ, Zhao Y, Sundin GW. Contribution of *Erwinia* amylovora Exopolysaccharides amylovoran and levan to biofilm formation: Implications in pathogenicity. Phytopathology. 2009;99(11):1237–44.

58. Tian Y, Zhao Y, Shi L, Cui Z, Hu B, Zhao Y. Type VI secretion systems of *Erwinia* amylovora contribute to bacterial competition, virulence, and exopolysaccharide production. Phytopathology. 2017;107(6):654–61.

59. Holub EB. Natural history of *Arabidopsis thaliana* and oomycete symbioses. Eur J Plant Pathol. 2008;122(1):91–109.

60. Kettle DS. A study of the association between moorland vegetation and breeding sites of Culicoides (Diptera, Ceratopogonidae). Bull Entomol Res. 1961;52(2):381–411.

61. Carpenter S, Mordue W, Mordue (Luntz) J. Selection of resting areas by emerging *Culicoides impunctatus* (Diptera: Ceratopogonidae) on downy birch (*Betula pubescens*). Int J Pest Manag. 2008;54(1):39–42.

62. Chen D, Shao Q, Yin L, Younis A, Zheng B. Polyamine function in plants: Metabolism, regulation on development, and roles in abiotic stress responses. Front Plant Sci. 2019;9:1945.

63. Michael AJ. Polyamine function in archaea and bacteria. Journal of Biological Chemistry. 2018;293(48):18693–701.

64. Choi O, Kang B, Lee Y, Lee Y, Kim J. Pantoea ananatis carotenoid production confers toxoflavin tolerance and is regulated by Hfq-controlled quorum sensing. MicrobiologyOpen. 2021;10(1):e1143.

65. Vijaya Kumar S, Abraham PE, Hurst GB, Chourey K, Bible AN, Hettich RL, et al. A carotenoiddeficient mutant of the plant-associated microbe Pantoea sp. YR343 displays an altered membrane proteome. Sci Rep. 2020;10(1):14985.

66. Bible AN, Fletcher SJ, Pelletier DA, Schadt CW, Jawdy SS, Weston DJ, et al. A carotenoid-deficient mutant in Pantoea sp. YR343, a bacteria isolated from the rhizosphere of Populus deltoides, is defective in root colonization. Front Microbiol. 2016;7:491.

67. Jia KP, Mi J, Ali S, Ohyanagi H, Moreno JC, Ablazov A, et al. An alternative, zeaxanthin epoxidaseindependent abscisic acid biosynthetic pathway in plants. Mol Plant. 2022;15(1):151–66.

68. Hirano T, Okubo M, Tsuda H, Yokoyama M, Hakamata W, Nishio T. Chitin heterodisaccharide, released from chitin by chitinase and chitin oligosaccharide deacetylase, enhances the chitin-metabolizing ability of *Vibrio* parahaemolyticus. J Bacteriol. 2019;201(20).

69. Hirano T, Aoki M, Kadokura K, Kumaki Y, Hakamata W, Oku T, et al. Heterodisaccharide 4-O-(N-acetyl-β-d-glucosaminyl)-d-glucosamine is an effective chemotactic attractant for *Vibrio* bacteria that produce chitin oligosaccharide deacetylase: *Vibrio* chemotaxis by disaccharides. Lett Appl Microbiol. 2011;53(2):161–6.

70. Shapiro LR, Scully ED, Straub TJ, Park J, Stephenson AG, Beattie GA, et al. Horizontal gene acquisitions, mobile element proliferation, and genome decay in the host-restricted plant pathogen *Erwinia tracheiphila*. Genome Biol Evol. 2016;8(3):649–64.

71. Leo JC, Oberhettinger P, Schütz M, Linke D. The inverse autotransporter family: Intimin, invasin and related proteins. Int. J. Med. Microbiol. 2015;305(2):276–82.

72. Hasson SO, Judi HK, Salih HH, Al-Khaykan A, Akrami S, Sabahi S, et al. Intimin (eae) and virulence membrane protein pagC genes are associated with biofilm formation and multidrug resistance in Escherichia coli and Salmonella enterica isolates from calves with diarrhea. BMC Res Notes. 2022;15(1):321.

73. Mundy R, Schüller S, Girard F, Fairbrother JM, Phillips AD, Frankel G. Functional studies of intimin in vivo and ex vivo: implications for host specificity and tissue tropism. Microbiology. 2007;153(4):959–67.

74. Facey PD, Méric G, Hitchings MD, Pachebat JA, Hegarty MJ, Chen X, et al. Draft genomes, phylogenetic reconstruction, and comparative genomics of two novel cohabiting bacterial symbionts isolated from Frankliniella occidentalis. Genome Biol Evol. 2015 Aug;7(8):2188–202.

75. Harada H, Oyaizu H, Kosako Y, Ishikawa H. *Erwinia* aphidicola, a new species isolated from pea aphid, Acyrthosiphon pisum. J Gen Appl Microbiol. 1997;43(6):349–54.

76. Campillo T, Luna E, Portier P, Fischer-Le Saux M, Lapitan N, Tisserat NA, et al. *Erwinia* iniecta sp. nov., isolated from Russian wheat aphid (Diuraphis noxia). Int. J. Syst. Evol. Microbiol. 2015;65(Pt_10):3625–33.

77. Chanbusarakum L, Ullman D. Characterization of bacterial symbionts in Frankliniella occidentalis (Pergande), Western flower thrips. J. Invertebr. Pathol. 2008;99(3):318–25.

78. Berry C, Meyer JM, Hoy MA, Heppner JB, Tinzaara W, Gold CS, et al. Biting Midges, Culicoides spp. (Diptera: Ceratopogonidae). Dordrecht: Springer Netherlands; 2008. p. 510–9.

79. Harada H, Ishikawa H. Experimental pathogenicity of *Erwinia* aphidicola to pea aphid, Acyrthosiphon pisum. J Gen Appl Microbiol. 1997;43(6):363–7.

80. De Vries EJ, Jacobs G, Sabelis MW, Menken SBJ, Breeuwer JAJ. Diet–dependent effects of gut bacteria on their insect host: the symbiosis of *Erwinia* sp. and western flower thrips. Proc R Soc Lond B. 2004;271(1553):2171–8.

81. Saldierna Guzmán JP, Reyes-Prieto M, Hart SC. Characterization of *Erwinia* gerundensis A4, an almond-derived plant growth-promoting endophyte. Front Microbiol 2021;12:687971

