## Supplementary figures and images for "Comparative genomics of a novel *Erwinia* species associated with the Highland midge (*Culicoides impunctatus*)"

### Supplementary File 1

#### Slide 1
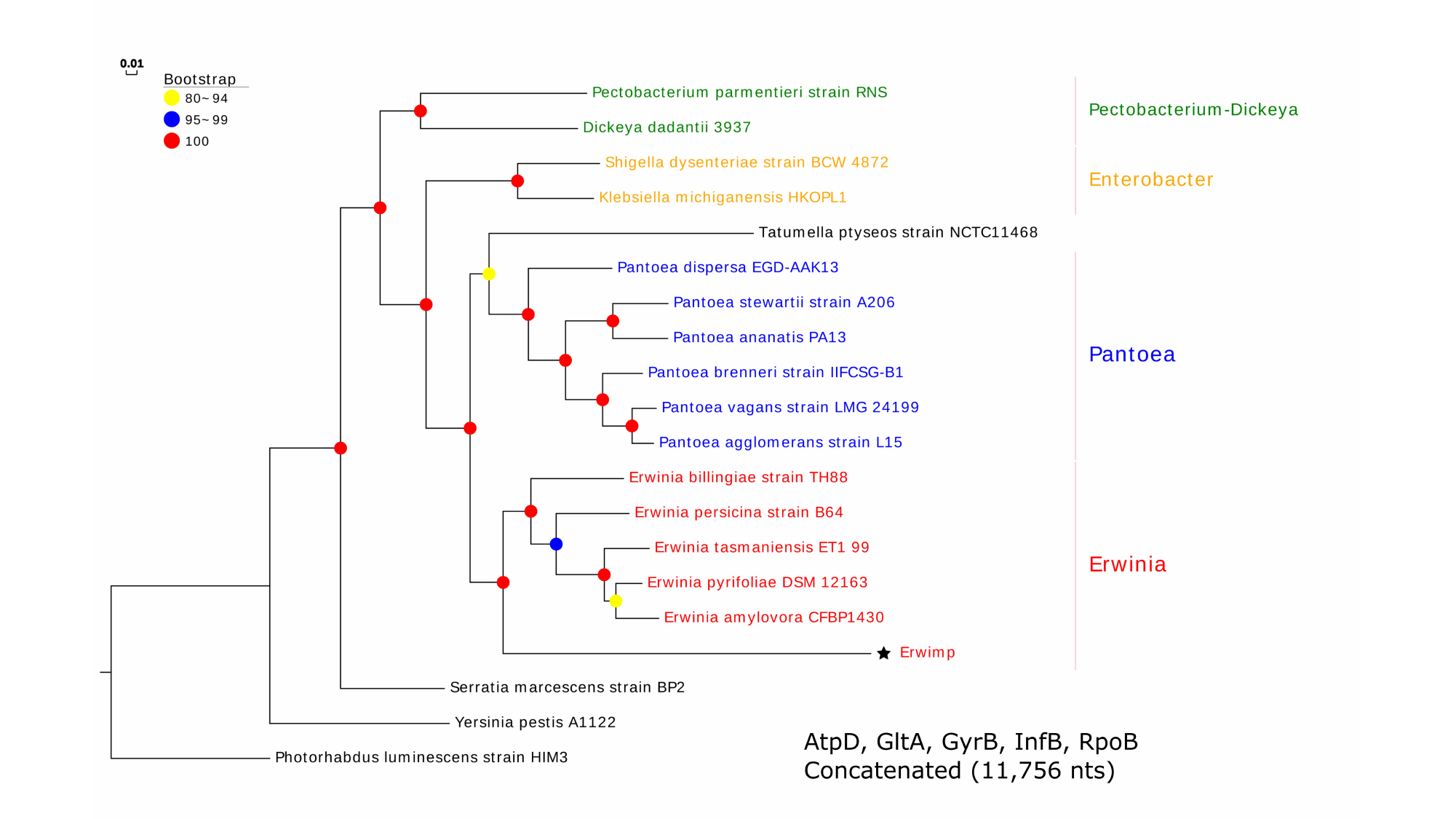
