## Supplementary File 6 for "Comparative genomics of a novel *Erwinia* species associated with the Highland midge (*Culicoides impunctatus*)"

### Slide 1
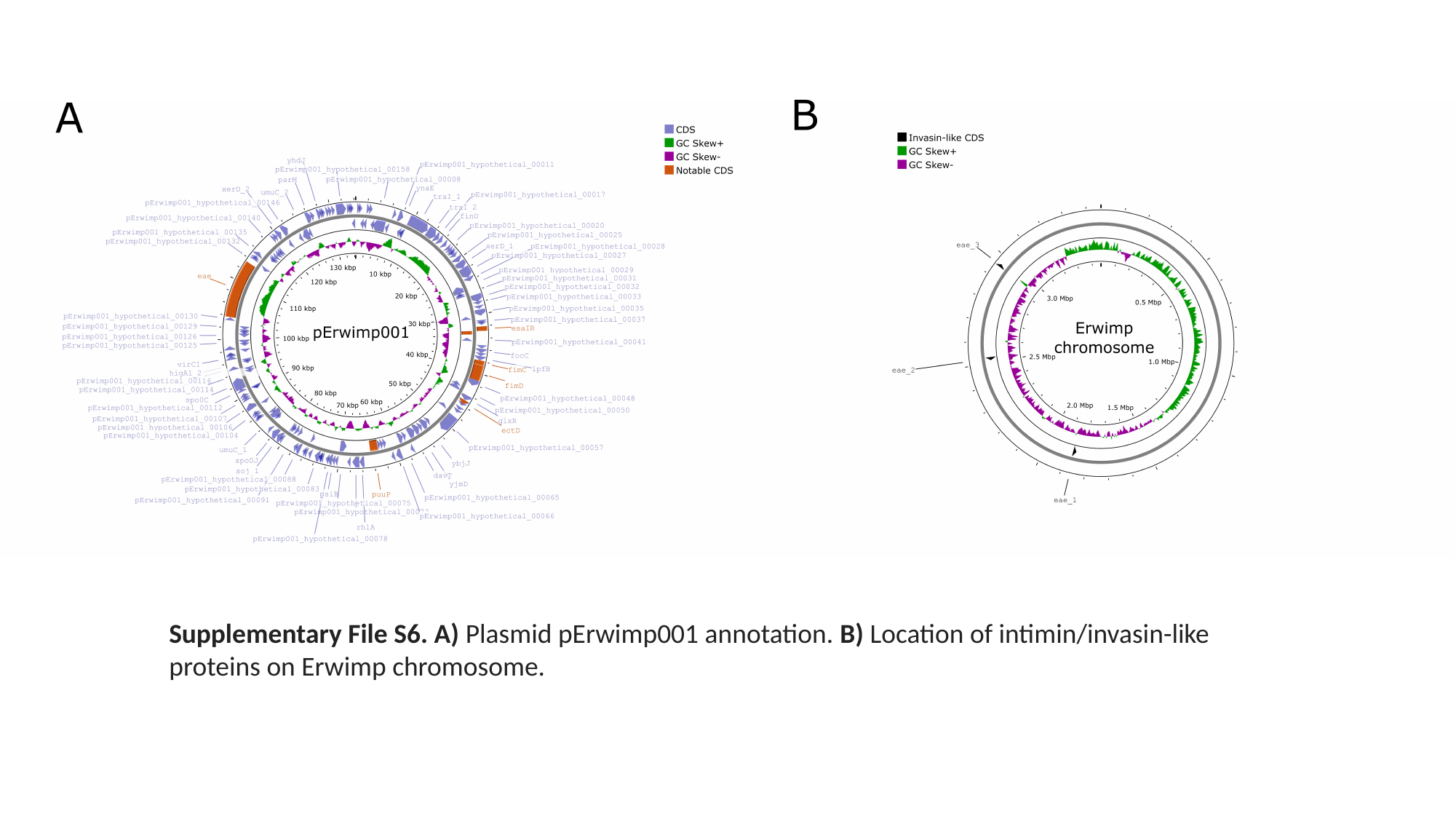

Supplementary File S6. A) Plasmid pErwimp001 annotation. B) Location of intimin/invasin-like proteins on Erwimp chromosome.
