## Supplementary File 4 for "Comparative genomics of a novel *Erwinia* species associated with the Highland midge (*Culicoides impunctatus*)"

# BUSCO version is: 5.1.2 
# The lineage dataset is: gammaproteobacteria_odb10 (Creation date: 2021-02-23, number of genomes: 976, number of BUSCOs: 366)
# Summarized benchmarking in BUSCO notation for file /var/www/gvolante/request/project/202309222236-1HD382EF5V1MY9WJ/Erwimp_chromosome_1.fa.ca
# BUSCO was run in mode: genome
# Gene predictor used: prodigal

	***** Results: *****

	C:99.7%[S:99.7%,D:0.0%],F:0.3%,M:0.0%,n:366	   
	365	Complete BUSCOs (C)			   
	365	Complete and single-copy BUSCOs (S)	   
	0	Complete and duplicated BUSCOs (D)	   
	1	Fragmented BUSCOs (F)			   
	0	Missing BUSCOs (M)			   
	366	Total BUSCO groups searched		   

Dependencies and versions:
	hmmsearch: 3.1
	prodigal: 2.6.3
